# PDX models of relapsed pediatric AML preserve global gene expression patterns and reveal therapeutic targets

**DOI:** 10.1101/2022.01.31.478534

**Authors:** Mark Wunderlich, Jing Chen, Christina Sexton, Nicole Manning, Luke Byerly, Eric O’Brien, John P. Perentesis, James C. Mulloy, Benjamin Mizukawa

**Affiliations:** Division of Experimental Hematology and Cancer Biology, Cancer and Blood Disease Institute, Cincinnati Children’s Hospital Medical Center, Cincinnati, Ohio, USA; Division of Biomedical Informatics, Cincinnati Children’s Hospital Medical Center, Cincinnati, Ohio, USA; Department of Pediatrics, University of Cincinnati College of Medicine, Cincinnati, Ohio, USA; Division of Oncology, Cancer and Blood Disease Institute, Cincinnati Children’s Hospital Medical Center, Cincinnati, Ohio, USA

**Keywords:** Patient Derived Xenograft, AML, RNAseq, KMT2A, CBFA2T3-GLIS2, CPX-351, venetoclax, NSGS

## Abstract

As patient-derived xenograft (PDX) models of acute myeloid leukemia (AML) become increasingly common tools for preclinical evaluation of targeted therapies it becomes important to consider the fidelity with which this system recapitulates the disease state found in patients. Gene expression profiling of patient blasts has been successfully used to identify distinct subtypes of AML to uncover sub-type specific vulnerabilities and to predict response to therapy and outcomes. Currently, there is little information regarding how well PDX models of AML mimic global gene expression patterns found in patients. In order to address this point, we performed detailed RNA-Seq analysis of data obtained from a diverse series of pediatric AML PDXs, separately and compared to primary patient data. When unsupervised clustering was applied to the PDX sample dataset, we found grouping associated with KMT2A (MLL) gene status. Additionally, in combined analysis, PDX samples were found to align with primary patient samples harboring similar genetics. We found a strong correlation of expression levels of nearly all expressed transcripts in PDX and patient datasets thus demonstrating faithful recapitulation of gene expression signatures. Furthermore, paired patient/PDX samples showed strong concordance, suggesting retention of sample-specific gene expression in immune deficient mice. Comparisons of PDX models propagated in NOD/SCID/IL2rg ^-/-^ (NSG) mice compared to NSG mice with transgenic expression of human SCF, GM-CSF, and IL-3 (NSGS) revealed minimal differences related to increased JAK/STAT and macrophage activation pathways in NSGS. Additionally, a unique RAM immunophenotype associated expression signature pointed to discovery of cryptic CBFA2T3-GLIS2 rearrangement as the mechanistic driver mutation in two PDX models. Based on the relatively high BCL2 mRNA in these models, we tested the efficacy of venetoclax in combination with CPX-351 which resulted in reduced leukemia burden and prolonged survival. These results validate the PDX system as surrogate of the molecular signatures in high-risk pediatric AML and highlight this system’s utility for pre-clinical therapeutic discovery, especially in very rare subtypes of disease.

## Introduction

PDX generation has become more commonplace in recent years as expectations grow that such systems will prove to be more relevant than mouse and cell line models and will therefore be more informative and predictive preclinical models of therapy^1-5^. Several groups have shown the ability of individualized PDX avatars to predict and even guide the personalized therapy of cancer patients^6-10^. However, the ability of PDX models to serve as faithful mouse avatars hinges on the retention of the specific molecular features of each case.

Many tumor types exhibit wide intra-tumor complexity and spatial distribution of distinct clones and metastasis, making personalized avatar modeling nearly impossible^11-13^. However, once established, there is evidence that PDX models are stable over many passages, allowing for a consistent, reliable system^4,14^. Despite this caveat, PDX models may still be good representatives of a specific disease subtype, if global gene expression patterns are retained.

Several studies have performed comparisons of various tumors and representative PDX models in order to determine how closely global gene expression in the xenograft setting reflects the patient condition. An NCI study found that PDX tumor gene expression largely mimicked the tumor type of origin in most samples tested, at least in so far as models clustered together by tumor type^15^. Importantly, PDX tumors were closer to primary tumors than were cell lines, suggesting a general benefit of in vivo propagation.

A recent study of 9 patient/PDX ovarian cancer pairs found ∽10% of genes were dysregulated in PDX, with most of those being downregulated in PDX and many attributable to the lack of immune system and human stroma cell interactions^16^. Similarly, 10 paired uveal melanoma/PDX models revealed a 3% difference in microarray gene expression results, the majority of which were related to decreased immune and stromal interactions and increased gene expression associated with cell cycle, kinase activity, and DNA repair in PDXs^17^. Similar differences were observed in 8 pancreatic cancer/SCID mouse PDX pairs^18^. On the other hand, a study of 20 colon cancer/PDX pairs found unstable genetic and expression profiles that changed over serial PDX passages and failed to recapitulate the subtype signature of the initiating patient^19^. This could be unique to the type of cancer or related to host strain, as this study utilized athymic nude mice for PDX generation and the relatively heightened level of residual immunity may pressure higher rates of evolution and subsequent diversion in these hosts. However, changes in growth characteristics and morphology have also been noted in PDXs of additional cancer types^20^. These results demonstrate the need for each PDX model system to be validated at the disease subtype and individual model levels.

Regarding leukemia, several studies have examined the fidelity of acute lymphoid leukemia (ALL) PDX model gene expression. A comparison of chip-based gene expression profiles and methylation patterns of 10 pediatric B-ALL patient/PDX pairs found high correlation of both profiles^21^. Similarly, pediatric T-ALL PDXs were found to retain 97.5% of the patient sample DNA methylation pattern^22^. Another microarray study using 5 matched pairs showed hierarchical clustering of 4 of 5 pairs while only 5% of transcripts varied in expression by 2-fold or more between patient and PDX^23^.

Furthermore, it has been demonstrated that ALL PDX models can harbor rare, dormant, niche-associated leukemic cells that have gene expression profiles resembling treatment resistant cells from patients with detectable minimal residual disease after therapy^24^, providing strong support for the use of leukemia PDX models to study preclinical therapeutic response.

In contrast to work done to validate gene expression of ALL PDX models, relatively little information is available for AML PDX models. One previous study that examined RNA sequencing data from 3 pediatric AML FLT3-ITD^+^ primary patient samples and matched PDX bone marrow samples found significant variation in gene expression levels at less than 5% of genes^25^. Another found that 95% of genes from a single AML sample (a dual FLT3-ITD/TKD mutant) were expressed at similar levels in primary and secondary NOD/SCID PDX^26^. A limited analysis of a set of 3 matched pediatric AML patient/PDX (FLT3-wt) found some degree of relatedness of gene array signatures by PCA analysis^27^. To date, there is a relative lack of a comprehensive analysis of RNAseq data from PDX models that broadly establishes strong retention of patient gene expression signatures for pediatric AML.

Here, we have performed RNA sequencing of a diverse panel of PDX models of pediatric AML in order to compare the global gene expression signatures to an independent patient sample database as well as matched original patient material. We also compared the effects of mouse strain on gene expression and found few consistent differences between parallel engrafted pairs of NSGS/NSG mice. Additionally, gene expression profiles enabled us to identify the CBFA2T3-GLIS2 (C/G) fusion present in two samples of our cohort which had previously unknown mechanisms of transformation. We then examined the unique C/G PDX signature to discover several potential avenues for improving the therapeutic response of these leukemias, including BCL2 inhibition, which resulted in reduced disease burden and longer survival when combined with chemotherapy in our preclinical PDX system.

Together, these data show that pediatric AML PDX models generally recapitulate global gene expression patterns found in patient samples and they can be used to generate hypotheses for preclinical testing of precision therapies.

## Materials and Methods

### Patient Samples

Patient BM and PB samples were obtained from residual diagnostic material from patients at CCHMC and used according to CCHMC institutional review board protocols #2008-0021 and #2010-0658. If specimens were not used immediately, aliquots were frozen in hetastarch freeze media (IMDM/5%BSA/5%DMSO/50%Hespan) after lysis of RBCs in ammonium chloride. Aliquots of some patient samples were frozen in RNALater solution and used for paired patient / PDX analysis.

### PDX

NSG, NRG, NSGS^28^, and NRGS^29^ mice were used as hosts. Primary mice were given a single 30mg/kg intraperitoneal dose of busulfan^30^ 24hrs prior to intravenous or intrafemoral injection of OKT3-coated cell preparations^29^. Serial transplants were performed by intravenous injection of approximately 1×10^6^ marrow or spleen PDX samples into recipients with (NSG/NRG) or without (NSGS/NRGS) prior busulfan pre-conditioning. Bone marrow aspirations were performed to monitor engraftment as described previously^31^. Within 30 minutes of sacrifice, the 4 long bones from the rear legs were crushed in 5mLs of cold PBS/3%FBS. 250uL of strained sample (5% total marrow) was centrifuged at 600g for 6 minutes and resuspended in RNA Lysis/Binding Buffer (mirVana miRNA kit, Thermo Fisher) and stored at −80C. AML engraftment percentages were determined from FACS analysis (FACSCanto II, BD) of preparations of BM and spleens at time of sacrifice. The FACS antibody panel consisted of mCD45-APCCy7 (30-F11), hCD45-FITC (HI30), CD3-PECy7 (SK7), CD13-PE (WM15), CD33-PE (WM53), CD19-VioBlue (LT19, Miltenyi Biotec), CD56-v510 (NCAM16.2), and CD34-APC (581). All were from BD unless noted. Anti-mouse and anti-human FcyR blocking antibodies were used to prevent non-specific binding (Miltenyi Biotec). 7-aminoactinomycin D (BD) was used to label dead cells for exclusion from analysis. Flow cytometry data was analyzed with FloJo software (BD).

### Total RNA extraction and QC

The RNA was extracted using the mirVana miRNA Isolation Kit (Thermo Fisher, Grand Island, NY) with the total RNA extraction protocol. In brief, cell lysates stored at −80C in Lysis/Binding Buffer were sent to the University of Cincinnati Genomics, Epigenomics and Sequencing Core for total RNA extraction according to the protocol and eluted with 60 µl Elution Buffer. The RNA concentration was measured by Nanodrop (Thermo Scientific, Wilmington, DE) and its integrity was determined by Bioanalyzer RNA 6000 Pico Kit (Agilent, Santa Clara, CA).

### Library preparation

NEBNext Poly(A) mRNA Magnetic Isolation Module (New England BioLabs, Ipswich, MA) was used for poly(A) RNA purification with 50 uL total RNA as input. SMARTer Apollo NGS library prep system (Takara, Mountain View, CA) was used for automated poly(A) RNA isolation. The library for RNA-Seq was prepared by using NEBNext Ultra II Directional RNA Library Prep kit (New England BioLabs, Ipswich, MA). During the second cDNA synthesis dUTP was incorporated to maintain strand specificity. In short, the isolated poly(A) RNA or rRNA/globin depleted RNA was fragmented (∽200 bp), reverse transcribed to the 1st strand cDNA, followed by 2nd strand cDNA synthesis labelled with dUTP. The cDNA was then end repaired and dA-tailed and ligated to the adapter. The dUTP-labelled 2nd strand cDNA was enzymatically removed to maintain strand specificity. After indexing via PCR (13 cycles) enrichment, the amplified libraries together with the negative control were cleaned up for QC analysis. To check the quality and yield of the purified library, one µL library was analyzed by Bioanalyzer (Agilent, Santa Clara, CA) using DNA high sensitivity chip. To accurately quantify the library concentration for the clustering, the library concentration was qPCR measured by NEBNext Library Quant Kit (New England BioLabs) using QuantStudio 5 Real-Time PCR Systems (Thermo Fisher, Waltham, MA).

### Sequencing

SR 1×51 bp sequencing: To study differential gene expression, individually indexed and compatible libraries were proportionally pooled (∽25 million reads per sample) for clustering in cBot system (Illumina, San Diego, CA). Libraries at the final concentration of 15 pM were clustered onto a single read (SR) flow cell v3 using Illumina TruSeq SR Cluster kit v3 and sequenced to 51 bp using TruSeq SBS kit v3 on Illumina HiSeq system.

### RNA-Seq analysis

Raw patient and PDX RNA Sequence reads were processed using the Illumina sequence analysis pipeline by the Statistical Genomics and Systems Biology Core at the University of Cincinnati. The pipeline used Tophat^32^ to align sequence reads in fastq files to human reference genome hg19 and generated bam files. The featureCounts function in the Rsubread package^33^ was used to create the counts table for genes. Samples that failed FastQC (http://www.bioinformatics.babraham.ac.uk/projects/fastqc) quality check or that contained low read counts or alignment rates were marked as low-quality samples and removed from further analysis.

In the bioinformatics analysis of the RNA-Seq data, a non-specific pre-filter was first applied to remove genes with low read counts. Genes with at least 0.5 reads per million (CPM) in at least N_s_ samples were kept for further analysis, where N_s_ is the number of samples in the smallest group. Principle component analysis (PCA) was used for unsupervised clustering of samples and the first two components were used to generate the plot for visualization. Differential expression analysis was performed by fitting linear regression models using the limma voom function^34^. For paired analysis, patient ID was included as the block effect. The p-values from the differential expression analysis was adjusted by False Discovery Rate (FDR). In all of our analysis, genes with FDR adjusted p-value < 0.1 were considered significant. Variance stabilizing transformed (VST) counts were created by the vst function in the DESeq2 package^35^ and used to perform PCA and hierarchical clustering. The heatmaps of differentially expressed genes were generated by the Morpheus package. The entire bioinformatics analysis was performed using the statistical computing platform R. Pathway over-representation analysis was performed using web application ToppGene^36^.

In some analyses we performed PCA and hierarchical clustering for PDX and TARGET samples together. In order to reduce the systematic difference caused by the two different data sources, the PDX and TARGET gene level counts were first VST transformed and per-gene normalized separately. In total, 11981 genes were found in common between the two datasets that passed our non-specific expression pre-filter.

### In vivo preclinical therapy model

Mice were engrafted with 1.5-2.3×10^6^ PDX spleen cells without conditioning by iv injection. Alternatively, 8×10^5^ M-07e AML cells (DSMZ, grown in RPMI/10%FBS with 10ng/mL IL-3) were injected into NSG mice without conditioning. Mice were randomly assigned to the treatment groups at day 14. Residual discard vials of CPX-351 (Vyxeos, Jazz Pharmaceuticals, 5mg/mL AraC and 2.2mg/mL daunorubicin) were obtained from the CCHMC pharmacy. It was aliquoted and stored at −20C. Immediately before use, vials were thawed and diluted 10-fold with PBS and administered to mice at a dose of 100uL per 10g body weight (measured before each dose) by iv injection for final doses of 5mg/kg AraC and 2.2mg/kg daunorubicin on day 1,3, and 5^37^. For traditional free daunorubicin and AraC combination therapy, we used the maximum tolerated dose which we previously determined to be 1.2mg/kg daunorubicin and 50mg/kg AraC on three consecutive days by iv injection^38^. Venetoclax (ABT-199, LC Laboratories) was dissolved in DMSO to 200mg/mL and aliquoted for storage at −20C in vials of 375uL. Each day one vial was thawed and 3.75mLs PEG400, 375uL Tween80, and 3mLs water were added, in order. Mice were dosed with 50uL per 10g body weight (oral gavage) for a final dose of 50mg/kg, Monday thru Friday for 4 weeks beginning at day 14.

## Results

### Minimal mouse read contamination in RNAseq analysis of pediatric PDX models with high engraftment

We began by producing RNAseq data from a series of PDX models developed in NSGS and NRGS mice from 26 unique patients (Table 1). These samples contained high levels of engraftment with most containing over 90% AML (range: 77-100%, mean: 94.1 +/- 5.9%). As a quality control check, we performed FastQ Screen^39^ analysis which attempts alignment of each read to both mouse and human genomes. The results showed low levels of mouse reads, in line with the levels of mouse cells and low level of reads matching both human and mouse genomes (Table 1, Fig 1A). In fact, a similar level of reads from pure patient samples (PS) also mapped to the mouse genome. The percentage of AML cells in the samples did correlate with alignment to human and mouse genomes (Fig 1B-C). Overall, we observed a read mapping rate of 61 +/- 6% with our PDX samples. These results suggested only very low contamination of the data with mouse reads, even without purification of human cells.

**Table 1.**
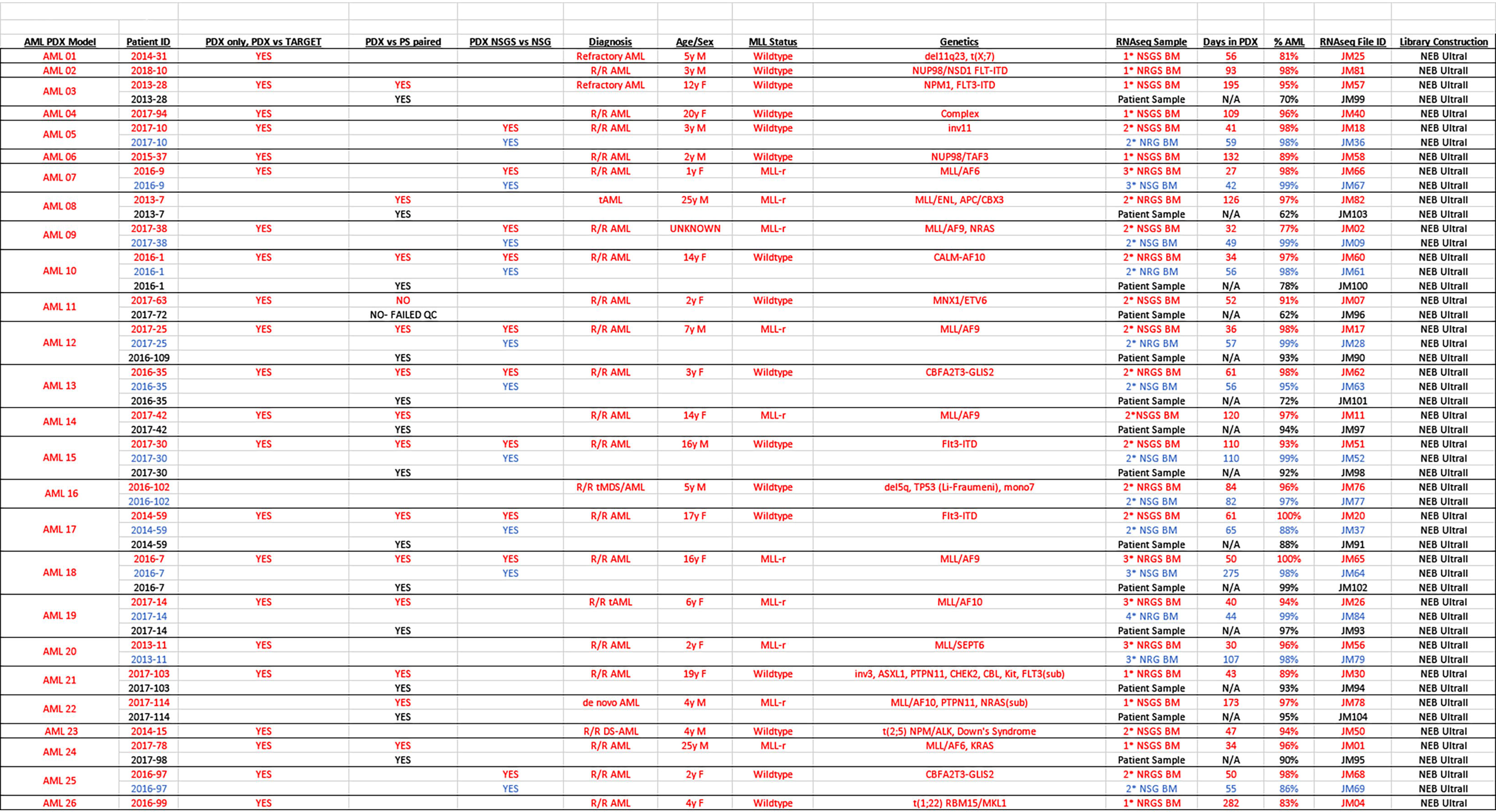
PDX model summary.

**Figure 1.**
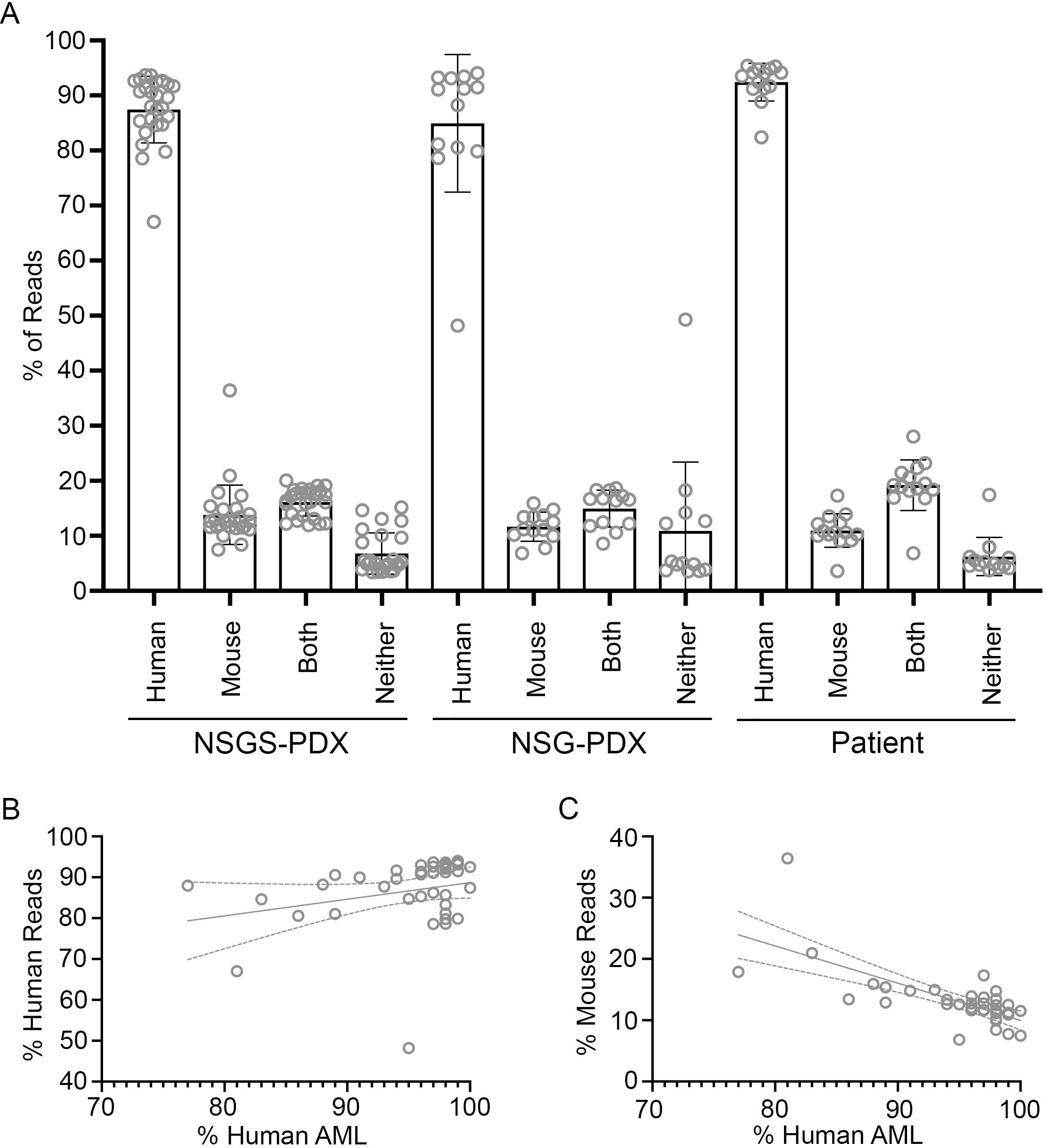
Alignment of RNA sequencing reads to human and mouse genomes. A) Bulk reads were aligned to human and mouse genomes with FastQ Screen. Percentages of all reads aligning to the indicated genomes are shown. B) Correlation of percentage of human reads with the percentage of human cells in the samples. C) Inverse correlation of percentage mouse reads with percentage of human cells.

### Host mouse strain has a limited influence on gene expression signatures

We set up our PDX models in NSG/NRG mice with transgenic expression of human myeloid-supportive cytokines SCF, GM-CSF, and IL-3 to promote the most efficient and robust engraftment of precious patient material^28^. To investigate any skewing effects that constitutive cytokine expression might have on gene expression, we also expanded 13 PDX models in NSG/NRG for sequencing. While the NSGS/NRGS models were significantly faster to harvest (NSGS/NRGS 50 +/- 24 days, NSG/NRG 81 +/- 62 days, p=0.0054 by Wilcoxon matched-pairs signed rank test), the samples consisted of similar levels of engraftment at sacrifice (NSGS/NRGS 95.5 +/- 5.9%, NSG/NRG 96.5 +/- 4.4%, p=0.4180) (Fig 2A-B, Table 1). Principle Component (PC) analysis revealed tight clustering of each NSGS/NSG pair (Fig 1C). We searched for genes with significantly altered expression levels between the NSGS/NRGS and NSG/NRG PDXs through a paired differential expression analysis which resulted in a list of 9 transcripts (Fig 2D-E). Consistent with the activities of the transgenic cytokines, 4 of these are well characterized members of the JAK/STAT pathway (SOCS1, GH1, CISH, and PIM1). Additionally, 3 others (MRC1, CLC, and HES1) are linked to aspects of macrophage activity and function^40-42^. This data suggests that transgenic SCF/GM-CSF/IL-3 cytokine expression did not exert consistent, widespread skewing of gene expression signatures.

**Figure 2.**
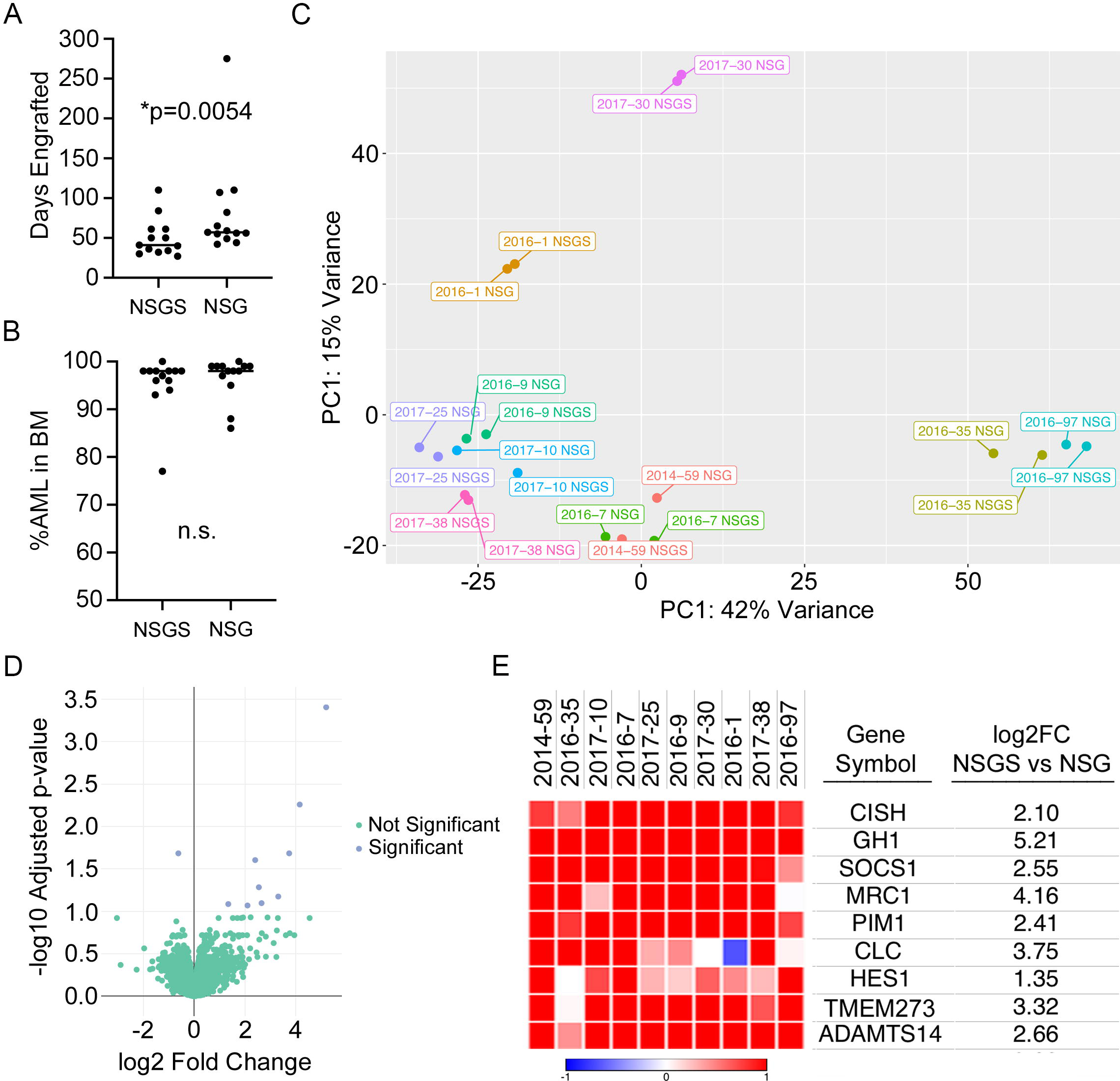
Comparison of NSGS and NSG PDX AML models. A) Latency of AML PDX models was significantly faster in NSGS mice, relative to NSG. B) Both NSGS and NSG mice developed AML with high percentage of human cells in the BM at sacrifice. C) Paired NSGS/NSG samples remain closely related by principal component analysis. D) Volcano plot of gene expression differences in NSGS models compared to paired NSG models. E) Heat map showing the 9 differential transcripts, all of which are higher in NSGS samples.

### PDX models of pediatric AML recapitulate global expression patterns associated with MLL-r

Our PDX cohort could be most evenly divided based on KMT2A (MLL) gene status, with 8 MLL rearranged (MLL-r) and 14 MLL wildtype (MLL-wt) samples. Therefore, we sought to identify MLL-r associated genes in our cohort. After filtering out low-expressed genes, 16853 genes were kept for further analysis.

PC analysis showed distinct separation of PDX samples based on MLL status (Fig 3A) with the first PC significantly associated with MLL status (Fisher’s exact test p-value 0.018). 228 genes were differentially expressed between MLL-r and MLL-wt (Fig 3B), 187 of which were found at higher levels in MLL-r samples while 41 were lower (Fig 3B, 3C, Table S1). Pathway analysis revealed that phosphatidylinositol binding, increased IgG level, and Immunoregulatory interactions between a Lymphoid and a non-Lymphoid cell were the most significant molecular function, mouse phenotype, and REACTOME pathway respectively (Table S2).

**Figure 3.**
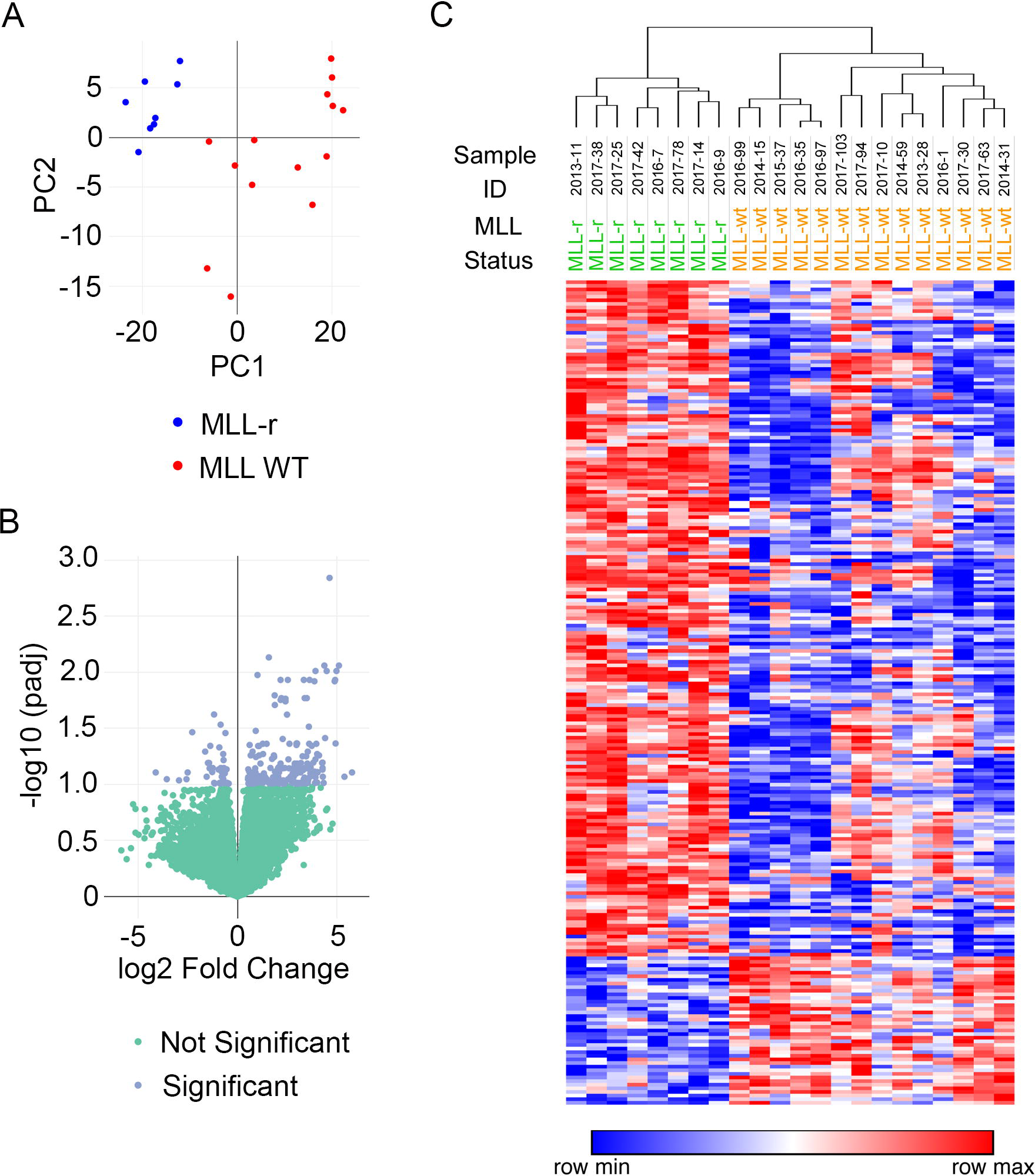
Generation of a PDX MLL-r signature. A) Principal component analysis, B) volcano plot, and C) heat map of NSGS PDX samples according to MLL status.

The samples that successfully generated PDX models in our hands were mainly from relapse and/or refractory cases. Notably, there were no PDXs from good prognosis categories with CBF or PML/RARA rearrangements. Considering the skewed composition of our cohort, we selected a similar set of pediatric AML PS RNA-Seq data from the NCI TARGET dataset. Due to a significant batch effect observed between the TARGET discovery and validation cohorts (data not shown), we limited the data to 46 samples from the discovery cohort (Table S3). After filtering out low-expressed genes, 14734 genes were kept for further analysis.

PCA analysis of the TARGET RNA-Seq dataset also showed a moderate grouping of 23 MLL-r and 23 MLL-wt samples (Fig 4A). The first PC was significantly associated with MLL status (Fisher’s exact test p-value 0.0072). We found 446 overexpressed and 382 under-expressed genes in the MLL-r cohort (Fig 4B, 4C, Table S4). A heatmap of differentially expressed genes showed several MLL-wt samples mixed into the MLL-r cluster and displaying portions of the MLL-r profile (Fig 4B).

**Figure 4.**
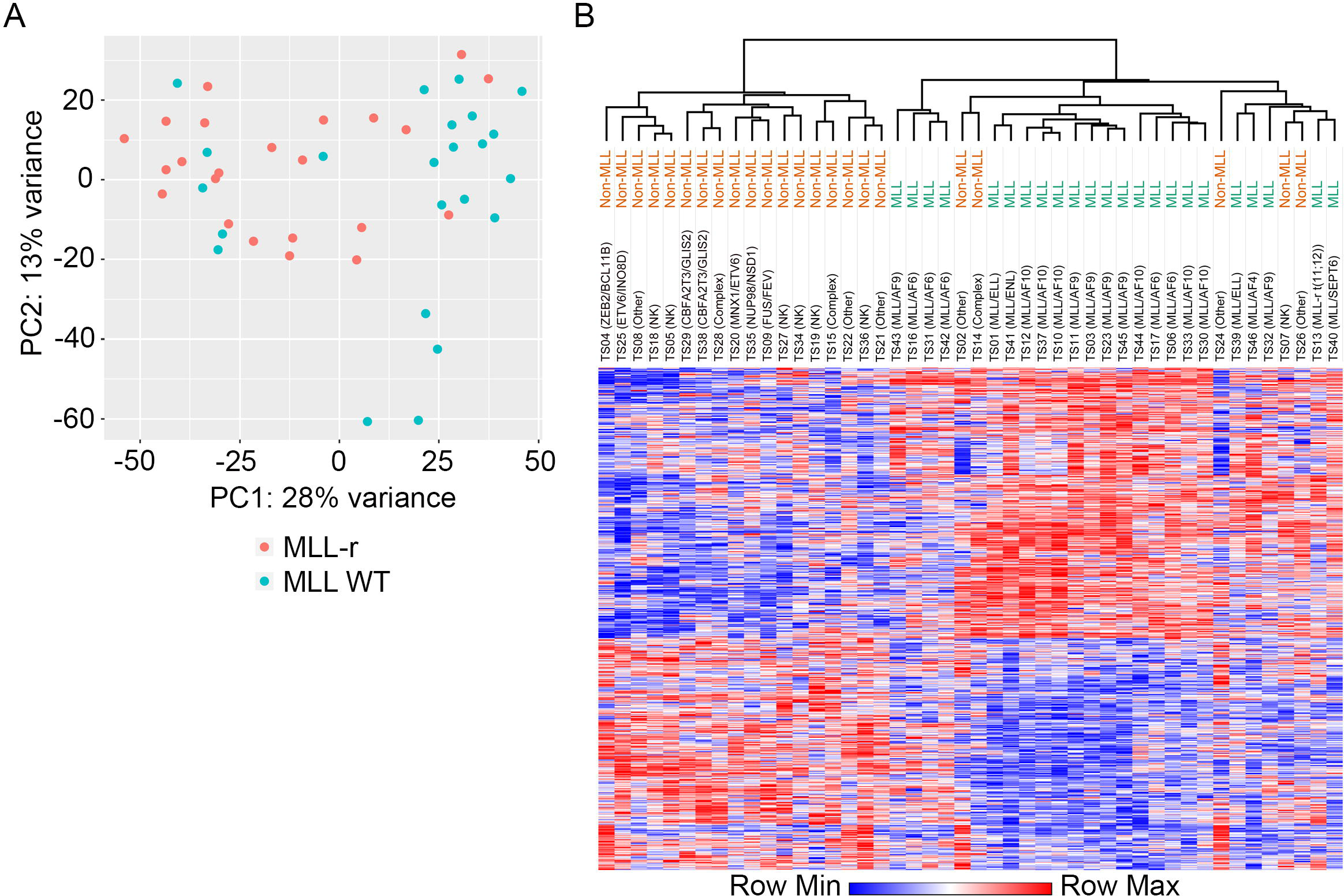
Generation of an MLL-r signature with NCI TARGET samples. A) Principal component analysis and B) heat map of selected TARGET patient sample RNA sequencing data according to MLL gene status.

Next, we reanalyzed combined PDX and TARGET data for PCA and hierarchical sample clustering based on 11913 common genes. Despite relatively small numbers and variable composition of MLL-r subtypes and other genomic aberrations in the MLL-wt groups, the PDX and TARGET results showed significant consistency. According to the PCA plot, MLL-r cases tended to cluster together, regardless of the source of material (Figure 5A). The first PC was significantly associated with MLL status (Fisher’s exact test p-value = 0.0039) but not the sample source (Fisher’s exact test p-value = 0.80). Similarly, the hierarchical clustering and heatmap (Figure 5B) of the 837 genes that were differentially expressed in either PDX or TARGET (190 in PDX, 702 in TARGET, 53 in both) showed that MLL-r specimens formed a distinct cluster. Notably, both degree and direction of differential expression in these 837 genes (as well as all expressed genes) was strikingly consistent between PDX and TARGET samples (Fig 5C, Table S5, correlation = 0.82, Fisher’s exact test p-value = 1.7e-139). Interestingly, two distinct MLL-r clusters were observed in the heatmap, with samples categorized as WHO M5 composing the larger group, while non-classical samples of various other WHO classifications dominated the second, smaller cluster.

**Figure 5.**
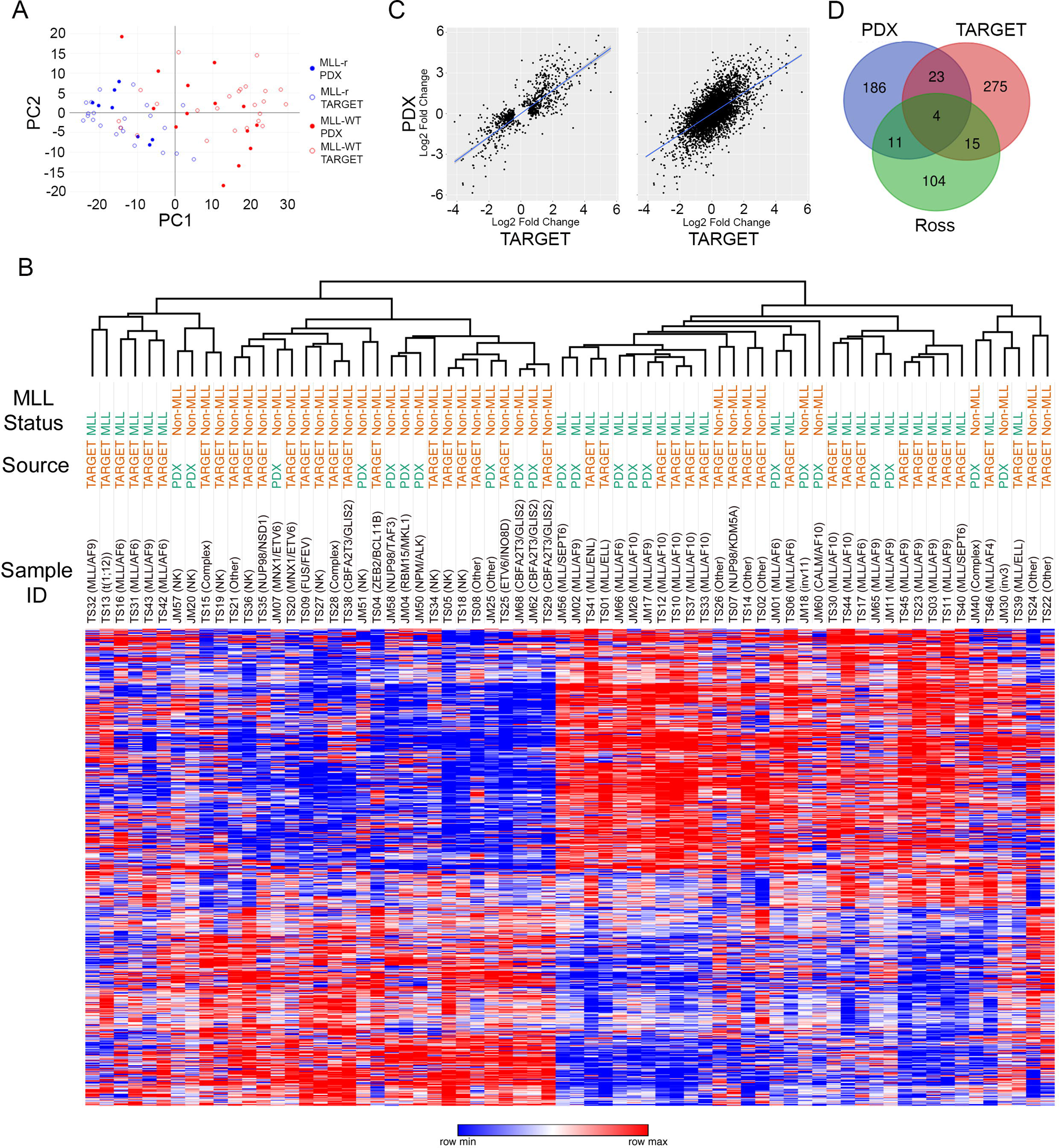
Generation of a mixed PDX/TARGET MLL signature. A) Principal component analysis of all samples with MLL status indicated. B) A heat map of all genes significantly correlated with MLL status. C) Correlation of expression between PDX and TARGET samples showing only MLL correlated genes (left) or all expressed genes (right). D) Venn diagram detailing the degree of overlap in identified MLL correlated genes between PDX, TARGET, or a separate study of primary patient material.

Additionally, we looked for overlap of the MLL-r PDX and TARGET gene lists with a previously published MLL class discriminating gene list from a large PS cohort^43^. While both of our gene lists resulted in similar overlap with the larger cohort (PDX 15/134, 11.2% and TARGET 19/134, 14.2%), each identified mostly unique genes with only 4 genes (APOC2, FEZ1, SAGE1, and TRPM4) present in all three gene lists (Fig 5D, Table 6).

### PDX models retain the original patient sample gene expression

Next, we compared 14 PDX models to matched PS for which available material with sufficient disease burden (>80%) was available. 2 samples were removed due to low sequencing quality. Despite selecting only samples with elevated levels of disease, the PDX models did contain significantly higher disease burden (PDX 95.9 +/- 3.2% vs PS 84.6 +/- 13.1%, p=0.0059 by Wilcoxon matched pairs rank test, Fig 6A). This is an important consideration because whereas the PDX are uniformly composed of clonal AML cells, the PS contains a variety of normal cells from myeloid and non-myeloid lineages which will alter bulk RNA composition. Because these PS and their PDX samples were processed in different batches, a direct paired test would likely be biased by the confounded batch effect. Therefore, we performed differential expression analysis separately in patient and PDX samples. Specifically, for each PS, we compared it with the other 11 samples to derive the ‘patient-specific’ gene expression signature and repeated this for PDX samples. We found that the PDX-derived patient-specific signatures significantly correlated with the patient-specific signatures derived directly from the PS, with mean correlation coefficient 0.54±0.14 (Table S6). 1535 genes were significant in at least one of the 24 signatures. PCA using these genes demonstrated that each PDX sample was strongly associated with the matched PS, although some degree of deviation was noted for some pairs (Fig 6B). These findings were reinforced by hierarchical clustering and heatmap (Fig 6C). We repeated hierarchical clustering based on all expressed transcripts, which again showed each PDX was closely related to its PS with one main outlier (Fig6D). Notably, this PDX was derived from a sample obtained at a later timepoint relative to the analyzed PS, after additional therapy with decitabine and vorinostat. Interestingly, after close examination, we found that the PDX lacked TP53 and IDH2 mutations that were observed in the PS, indicating that a distinct subclone was selected either by epigenetic therapy or by the xenograft procedure. The PS clustered near another sample with TP53 mutation and away from other MLL-r samples, indicating that the TP53 mutation likely results in strong global gene expression changes and highlighting the need for complete characterization of each PDX model.

**Figure 6.**
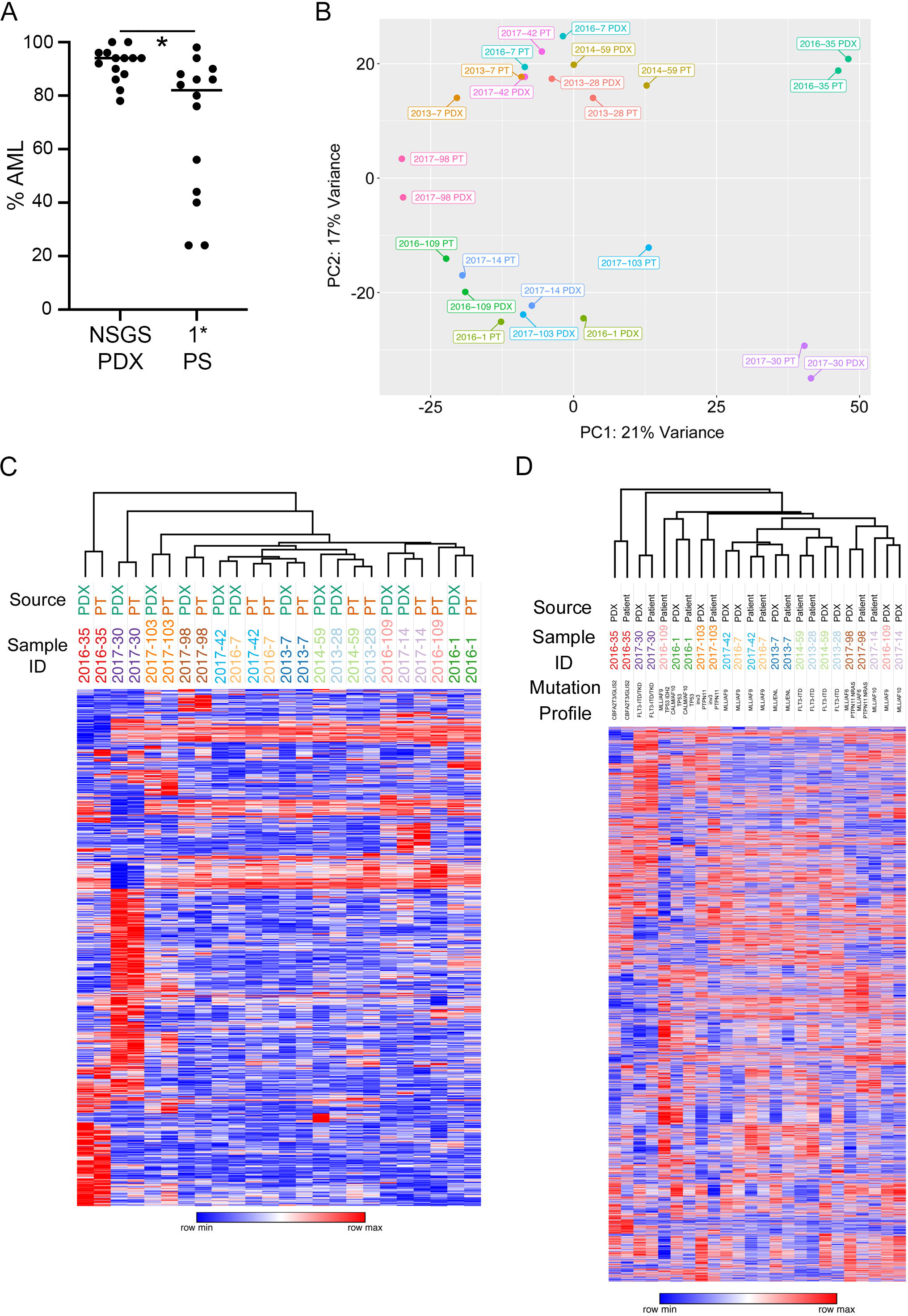
Paired PDX/Patient Sample analysis. A) Comparison of the leukemic percentage of the BM samples used. For PDX, percentage was determined by %hCD45+CD33+ cells at sacrifice. For patient samples, clinically reported blast percentages were used. B) PCA plot with paired samples in the same color. C) Each individual patient sample was compared to all others in order to generate a list of patient specific genes. This process was repeated for each sample and the combined list was used for hierarchical clustering. D) Alternatively, hierarchical clustering was generated using all expressed genes.

### PDX RNA-Seq allows for target discovery for rare subtype AMLs

In our paired NSG/NSGS analysis, there were 2 sets of samples that appeared similar to each other and quite distinct from all others (Fig 2C). Interestingly, they were both from a rare subtype of pediatric AML previously described as RAM immunophenotype which is based on surface expression of CD45, CD34, CD38, CD56, and HLA-DR and correlated with extremely poor prognosis^44^. At the time these samples were acquired, little was known about the driver mutations for RAM AML and clinical sequencing of these samples did not reveal any candidates.

To gain some insight into the biology of these samples, we reanalyzed the dataset and found a robust RAM-associated profile (Table S7). PC analysis resulted in these two samples clustering together with a subset of AMKL (Fig 7A). One of our samples was clinically classified as AMKL while the other was not. The most notable gene on this list of top 20 overexpressed genes is GLIS2, known for its involvement in CBFA2T3-GLIS2 (C/G) rearranged pediatric AMKL (Fig 7B). Remarkably, 16 of the top 20 (80%) overexpressed RAM phenotype genes were also found significantly upregulated by C/G expressing pediatric AMKLs^45^. In total, our results identified 341 of the 894 genes (38.14%) in the complete list. This finding along with the recent demonstration of overlap between RAM phenotype and C/G AMLs strongly suggested the presence of C/G in our two RAM samples^43^. We performed fusion gene transcript detection with JAFFA software^44^ and RT-PCR with C/G specific primers^45^, both of which confirmed CBFA2T3-GLIS2 rearrangement (Fig 7C and data not shown). Our C/G+ samples exhibited strong gene expression correlation with C/G+ samples found in the TARGET dataset (Fig 7D).

**Figure 7.**
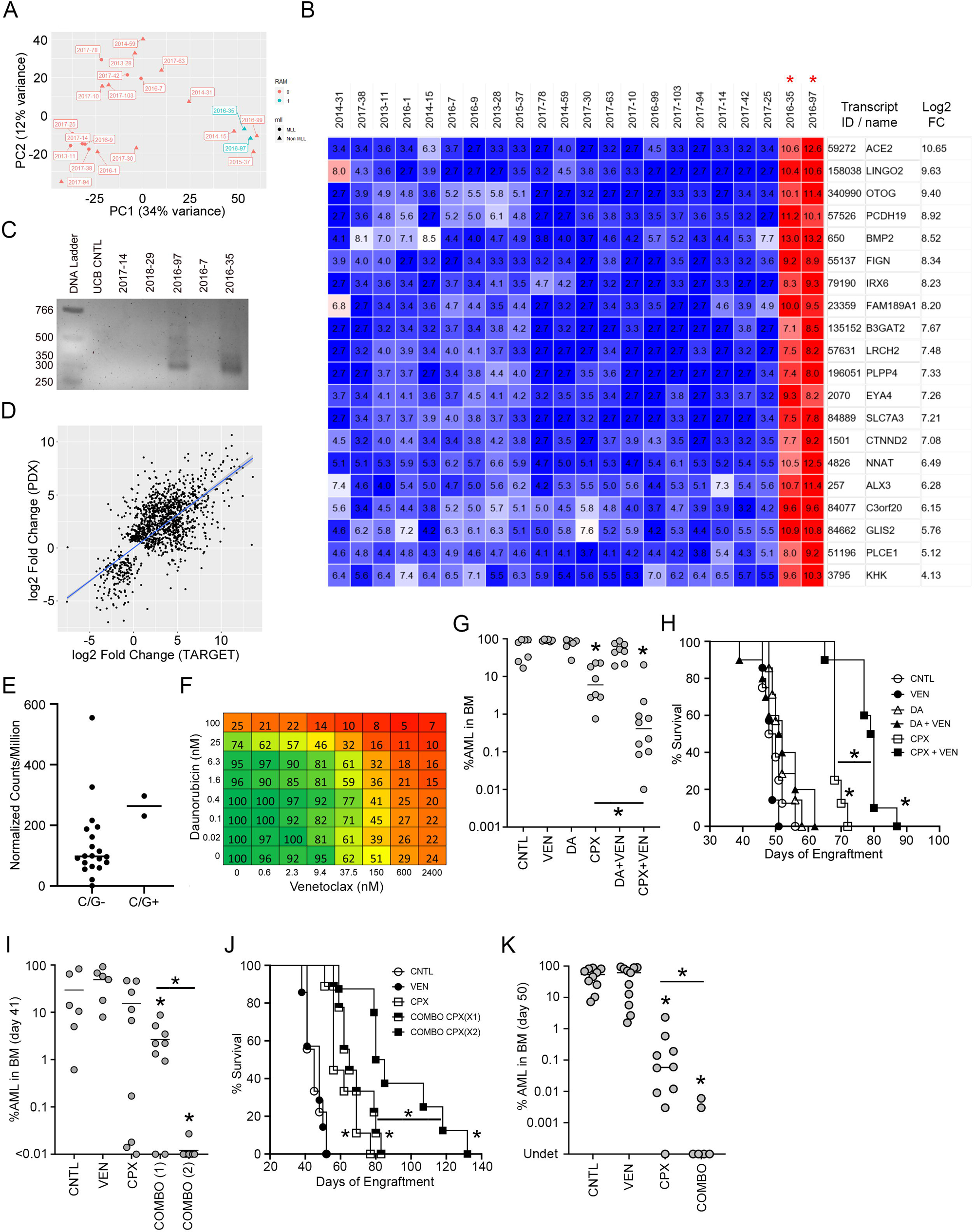
Demonstration of RNAseq-enabled discovery of a disease-specifc treatment option. A) PCA analysis of NSGS vs NSG PDX models shows distinct clustering of 2 samples with unknown genetics (blue font) together with AMKLs. B) The 2 PDX with unknown driver mutations were compared to all other PDX models. The heat map shows the top 20 overexpressed genes. C) RT-PCR detection of the CBFA2T3-GLIS2 (C/G) fusion gene in both models with unknown driver mutations. D) Comparison of gene expression signatures in C/G+ samples from PDX and TARGET datasets. C/G+ samples were compared to C/G- samples from the same dataset. Then the fold change for PDX and TARGET C/G+ samples were plotted against each other. E) BCL2 gene expression from PDX RNAseq data for C/G+ samples. F) MTT assay showing synergistic effects of combined daunorubicin and venetoclax exposure. The heat map numbers represent the viable cell percentage relative to vehicle control wells. G) NSGS mice were engrafted with PDX 2016-35 and subjected to the indicated therapy. BM aspirates were taken to assess disease burden on day 45 (32 days post therapy initiation). CNTL = control, VEN = venetoclax, DA = daunorubicin + Ara-C, CPX = CPX-351, a liposomal daunorubicin + Ara-C formulation. H) Survival of the mice in G. I) Day 42 disease burden (28 days post therapy) and J) survival of a second C/G+ PDX, 2016-97, in NRGS mice. Combo = combined CPX-351 and venetoclax with either 1 or 2 cycles of CPX-351. K) Day 50 BM disease burden for an NSG M-07e cell line xenograft experiment. Mice were treated for 1 week and marrow was assessed on day 32 following the start of therapy.

Our C/G gene list suggested several targetable pathways, including the extracellular matrix interaction, neurotrophin signaling, connexin and gap junction function, and BCL apoptotic pathways. While increased BCL2 expression has long been observed in R/R AMLs^46,47^, our C/G samples contained some of the highest levels compared to the other samples in our PDX cohort (Fig 7E) which we believed might predict exceptional sensitivity to venetoclax (ABT-199), a BCL2 specific inhibitor approved for use in older adults with chronic lymphoid leukemia or AML in combination with azacytidine, decitabine, or low dose cytarabine (AraC).

In vitro drug incubation assays with venetoclax and daunorubicin showed synergistic activity against a C/G+ ex vivo PDX sample (Fig 7F). We next performed an in vivo therapy experiment with NSGS mice engrafted with this PDX followed by treatment with standard induction chemotherapy (daunorubicin + AraC, DA), CPX-351 (Vyxeos, liposomal daunorubicin + AraC), venetoclax, and combinations of venetoclax with either formulation of induction chemotherapy. As with controls, mice that received venetoclax, DA, or DA + venetoclax had extremely high levels of AML in the BM after the 4-week treatment course (Fig 7G). However, CPX-351 treated mice had significantly reduced levels, consistent with reports of improved efficacy of this formulation^37,48^. Additionally, the CPX-351 + venetoclax group had a significant further reduction in AML, with 3 of 10 mice falling below the clinically relevant flow cytometry based MRD threshold of 0.1%. Importantly, this decreased disease burden translated into an increased survival advantage for the mice that received CPX-351 and venetoclax (Fig 7H). We tested the combination of venetoclax with CPX-351 in a second C-G PDX model and found that the combination was required to significantly decrease post therapy disease burden (Fig 7I). An extra cycle of CPX-351 during the venetoclax treatment window significantly enhanced both response to therapy (8 of 8 mice were MRD-) and survival (Fig 7I-J). A third xenograft model utilizing the M-07e cell line which is also C/G+^49^ also showed improved response to combination therapy with all mice below MRD and 4 of 6 with undetectable engraftment (Fig 7K).

## Discussion

The use of PDX-expanded samples for RNA-Seq analysis results in several benefits in the research setting. First, this approach allows for a more standardized workflow in which fresh and viable cells are obtained and processed uniformly. PS are often not immediately available for research and can only be processed only after lengthy delay, even when samples are guided to a central biorepository. Post therapy pediatric samples often lack enough material for research at all, which means we must rely on minute quantities of discarded diagnostic specimens that become available several days after collection. Additionally, xenograft leukemic expansion effectively outcompetes normal HSC engraftment (especially in serial transplants) and therefore eliminates the “contaminating” signal from non-malignant lymphocytes. PDX expansion results in samples with high blast content, generally over 90%. Pediatric PS available for research often consists of scant cellularity and may contain only low to MRD levels of disease, as is often the case with post therapy or relapse/refractory samples. These can often be expanded by PDX to acceptable levels. Samples are also at different timepoints of therapy, or even mid-therapy, which is a confounding factor that greatly affects gene expression. Another benefit to building such a PDX database is that we focus on studying only the samples that successfully engraft mice, a trait which is correlated with worse prognosis^50-52^ and patient survival^52,53^. Importantly, observations and hypotheses stemming from data analysis can be followed up experimentally in a sample specific manner.

We did not sort our PDX BM samples. Previous studies using the Affymetrix gene expression hybridization platform showed that low levels of contaminating mouse cells did not alter the quantification of the human profile^54^. A study of paired ovarian cancer patient-PDX RNA-Seq data showed that approximately 1% of reads could be aligned to either mouse or human genomes^16^. However, with RNA sequencing, we found significantly more overlap between human and mouse reads in our samples, but that level was similar with PS without any potential for mouse cell contamination, indicating a low likelihood for skewing effects. This finding is in line with that reported by others working with RNAseq data^55^.

Additionally, we did not observe widespread differences between PDX models generated in NSGS compared to paired NSG mice, with differences mainly related to JAK/STAT pathway activation that would be expected from constitutive overexpression of SCF, IL-3, and GM-CSF. This finding implies that the in vivo gene expression profiles for individual AMLs is driven by cell-intrinsic factors, and this view is bolstered by our observation of high fidelity between matched patient/PDX pairs. The use of NSGS mice may be preferred for many therapy models, since they allow for the faster expansion of many AMLs and can successfully engraft some samples that fail to engraft NSG. However, it is possible that therapies that target the JAK/STAT pathway could be blunted by the transgenic cytokine expression in NSGS and this should be considered. In fact, it has been shown that GM-CSF and IL-3 expression in NSGS mice accounts for a JAK2/STAT5-dependent mechanism of resistance to FLT3 inhibitors not observed in NSG mice^56^. One might argue that this makes NSGS the more relevant model, considering patient blasts are exposed to these signals in the human bone marrow microenvironment. However, cytokine expressing mice have been shown to promote solid tumor formation from AML cell lines, therefore they are often not ideally suited for all CDX applications^57^.

The rarity of many pediatric AML genetic subtypes makes performing clinical trials to establish the efficacy of personalized therapies in a timely manner quite challenging. Development and identification of candidate therapies for clinical trials outpaces the number of patients that can be recruited. Considering the rarity of pediatric leukemia, this holds true for even the most common subtypes of AML. It is possible that cohorts of PDX avatars could potentially stand in for patients in preclinical trials to narrow the field, provided proper validation is done prior to the undertaking in order to increase confidence that only the best potential therapies and dosing schedules and combinations are forwarded to real world testing. Confirmation of PDX retention of global RNA gene expression patterns is one important piece of that validation.

The observation that paired PS/PDX RNAseq samples clustered together is compelling evidence that the mouse microenvironment does not dramatically skew the global gene expression profiles. This could not have been simply assumed, as several studies of solid tumors found significant alterations in PDX models, presumably from differences in mouse vs human stromal interactions and effects of immune activity vs growth in an immune deficient host. We did not observe consistent differences in these genes, which may indicate fundamental differences in blood vs solid cancers. One explanation could be related to the relatively simplified clonal architecture in AML. It has been shown that AML (and especially pediatric AML) has one of the lowest mutation rates among many common cancer types and few clones predicted by bulk whole exome sequencing or karyotype analysis^58^. Mutation rate is also lower in pediatric AMLs with higher risk genetics, such as those in our cohort (MLL, NUP98, or GLIS2 rearrangement), compared to good prognosis samples that only rarely generate PDX models (such as those with CBF aberrations)^58,59^.

Our PDX cohort contained 2 samples from a rare subtype of pediatric AML, known as RAM phenotype. This high-risk group was identified as being 2.3% of patients from the Children’s Oncology Group clinical trial AAML0531 that exhibited very bright CD56, negative/dim CD45 and CD38, and negative staining for HLA-DR. RAM patients were more likely to be younger and the phenotype was associated with higher incidence of M7 AMKL classification. Interestingly, our RNA-Seq analysis of the 2 RAM phenotype samples showed that GLIS2 was one of the most highly overexpressed transcripts. A cryptic CBFA2T3-GLIS2 transcript was identified in pediatric AMKLs and defined a poor prognosis group^45^. Another study found the fusion in approximately 8% of cytogenetically normal pediatric AMLs, half of which were AMKL and half of which were other FAB sub-types^60^. A COG study found that 10 of 16 RAM phenotype samples harbored the CBFA2T3-GLIS2 rearrangement^61^. Expression of either CBFA2T3-GLIS2 or GLIS2 in murine progenitors led to enhanced self-renewal as demonstrated by serial methylcellulose replating^45,62^. The fusion was not previously detected in our samples (1 AMKL, 1 M0 with minimal differentiation), but our list showed remarkable correlation to a gene list generated by comparing CBFA2T3-GLIS2+ AMKL to other AMKLs^45^. CBFA2T3-GLIS2 AMKLs demonstrated altered SHH, WNT, and BMP pathway signaling (particularly BMP2 and BMP4 overexpression)^45^. CBFA2T3-GLIS2 transduced HEL cells showed an upregulation of ERG and downregulation of GATA1 which was similar to that observed in PS^62^. This system also produced differential gene expression which correlated well with patient data (r=0.48, p=0.017).

RAM immunophenotypes respond poorly to conventional DA induction therapy. While standard DA therapy produced no benefit in our PDX model, a liposomal formulation, CPX-351 (Vyxeos), reduced disease burden and prolonged lifespan. This finding is consistent with the finding of improved response, including overall survival, in adults with de novo AML^63^ or relapsed poor prognosis AML^64^ treated with CPX-351 compared to standard AraC and anthracycline regimens. Our RNAseq data showed increased BCL2 expression in C/G samples, even in comparison to many MLL-r AMLs which are known to upregulate this survival pathway. This finding led us to hypothesize that C/G AMLs might display enhanced sensitivity to venetoclax. In our in vivo preclinical PDX model, the CPX-351/venetoclax combination proved more effective than CPX-351 alone, with several mice measured as MRD- after therapy. MRD- status prior to BMT correlates with improved outcomes and BMT may be an approach to improve outcomes for pediatric AML with the RAM immunophenotype^65^.

We decided to focus initially on inhibition of BCL2 with venetoclax, as there are at least 4 ongoing clincial trials to investigate CPX-351/Venetoclax combination for a variety of AML diagnoses in the United States (clinicaltrials.gov, search results 4/6/20) which makes these results directly translatable. However, our RNAseq data contains other potential pathways that could be directly targeted in our system. For example, the GO neurotrophin signaling pathway and neurotrophin TRK receptor signaling pathway gene lists were found to be upregulated in our C/G samples. A previous report has demonstrated that CBF AMLs have increased levels of NTRKA protein (the TRK1 product) and can respond to NGF with increased proliferation^66^. While 44% of AMLs had detectable expression of TRK1, NGF (the ligand for NTRKA) was not detected in any tested PS^67^. Our C/G samples had upregulation of both NGF and TRK1 transcripts, suggesting a possible autocrine feedback loop. Such a feedback loop induced by dual overexpression of NGF/NTRKA has been demonstrated to protect 32D cells from apoptosis and lead to full leukemic transformation^68^. A constitutively active NTRKA isolated from a patient with AML had similar transforming effects^69^. It will be interesting to make use of our PDX C/G models to also test the efficacy of FDA approved drugs larotrectinib and entrectinib which have recently been approved for TRK-rearranged and mutated cancers.

RNAseq pathway analysis also uncovered upregulated gap junction components and connexin pathway members. AML cells engaged with stroma cells are more likely to be quiescent and protected from chemotherapy induced apoptosis in a gap junction dependent manner^70^. One potential mechanism for this phenomenon is through the transfer of stroma derived mitochondria, which has been shown to occur in AML PDX models in NSG mice and leads to resistance to standard AML induction agents^71^. Recently, carbenoxolone, an inhibitor of gap junction intercellular communication, has been shown to enhance apoptosis induced by AraC in cell lines and patient blasts in in vitro stroma co-cultures^72^. It will be interesting to test this approach with CPX-351+/- venetoclax in our PDX models to attempt to further improve treatment response and more eliminate MRD more consistently.

This demonstration with RAM phenotype / CBFA2T3-GLIS2 rearranged samples suggests that potential insights into other rare and distinct pediatric leukemia subtypes such as Pure Erythrocyte Leukemia (PEL) or Blastic Plasmacytoid Dendritic Cell Neoplasm (BPDCN) could be made with this approach. A study of 25 BPDCN PS which included 4 pediatric cases found a distinct signature compared to the closely related normal myeloid-derived resting pDCs^73^. Gene expression analysis of BPDCN vs normal pDC cells suggested NF-kB as a potential therapeutic target with bortezomib showing some in vitro effects on a cell line. Another study of a smaller cohort compared BPDCN to cutaneous AML and noted upregulated FLT3 which could potentially be targeted for therapy^74^. Similarly, PEL is another rare and aggressive subtype which has been difficult to classify, despite its unique morphology and immunophenotype^75^. The PDX RNAseq approach will afford us the opportunity to compare these rare leukemias to other aggressive AMLs in our dataset to attempt to verify these or uncover additional novel pediatric-specific targetable mechanisms that could be directly tested in the same preclinical PDX models.

## Author Contributions

MW conceived the project, designed experiments, performed experiments, analyzed data, interpreted results, and wrote the manuscript. JC contributed to the study design, designed and performed experiments, analyzed and interpreted data, and wrote the manuscript. EO designed and performed experiments, analyzed and interpreted data, and supplied patient samples. NM performed experiments and analyzed data. CS performed experiments. LB provided patient samples and patient RNA. JPP provided patient samples, funding, and other support. JCM and BM contributed to the study design, interpreted results, and edited the manuscript. JCM also contributed funding. All authors reviewed, edited, and approved of the manuscript.

## Acknowledgements

This work was supported by NIH/NCI grants R50 #CA211404 (MW), R01 #CA204895 (JCM), and R01 #CA215504 (JCM) and a CCHMC ARC award (JPP and JCM).

